# Deletion of FAT1 in hybrid EMT cells stimulates the migration of neighboring non-mutant cells through secretion of extracellular vesicles

**DOI:** 10.1101/2023.09.06.556588

**Authors:** Juliana Geay, Shailaja Seetharaman, Benoît Vianay, Matthieu Gélin, Olivia Fresnoy, Laurent Blanchoin, Manuel Théry

## Abstract

Cells in hybrid state of the epithelial-to-mesenchymal transition (EMT) have been shown to be responsible for tumor cell metastasis. However, the precise mechanisms underlying the morphological changes and acquisition of invasive phenotypes in hybrid EMT cells are still unknown. Here, we introduced the deletion of a proto-cadherin and well described oncogene, FAT1, in skin carcinoma cells to generate a hybrid state of EMT. Surprisingly, the FAT1 knock-out (KO) cells were less motile than the parental non-mutated cell line they were derived from. However, we observed that FAT1 KO cells secrete specific factors in the form of extra-cellular vesicles into their microenvironment, which promote the migration of surrounding non-mutant cells. When stimulated with these extracellular vesicles, groups of non-mutated parental cells collectively migrated faster and formed finger-like instabilities at the migrating front. Furthermore, we found that the actomyosin contractility of FAT1 KO cells in hybrid EMT states was much lower than the parental cells. It appeared that the factors secreted by FAT1 KO cells relaxed the traction forces in recipient cells. This force release likely fostered the scattering and migration of non-mutated cells surrounding FAT1 mutant cells. Thus, we characterized a non-autonomous promotion of cell invasiveness in the cancer cells surrounding FAT1-deficient cells.

**Figure:**
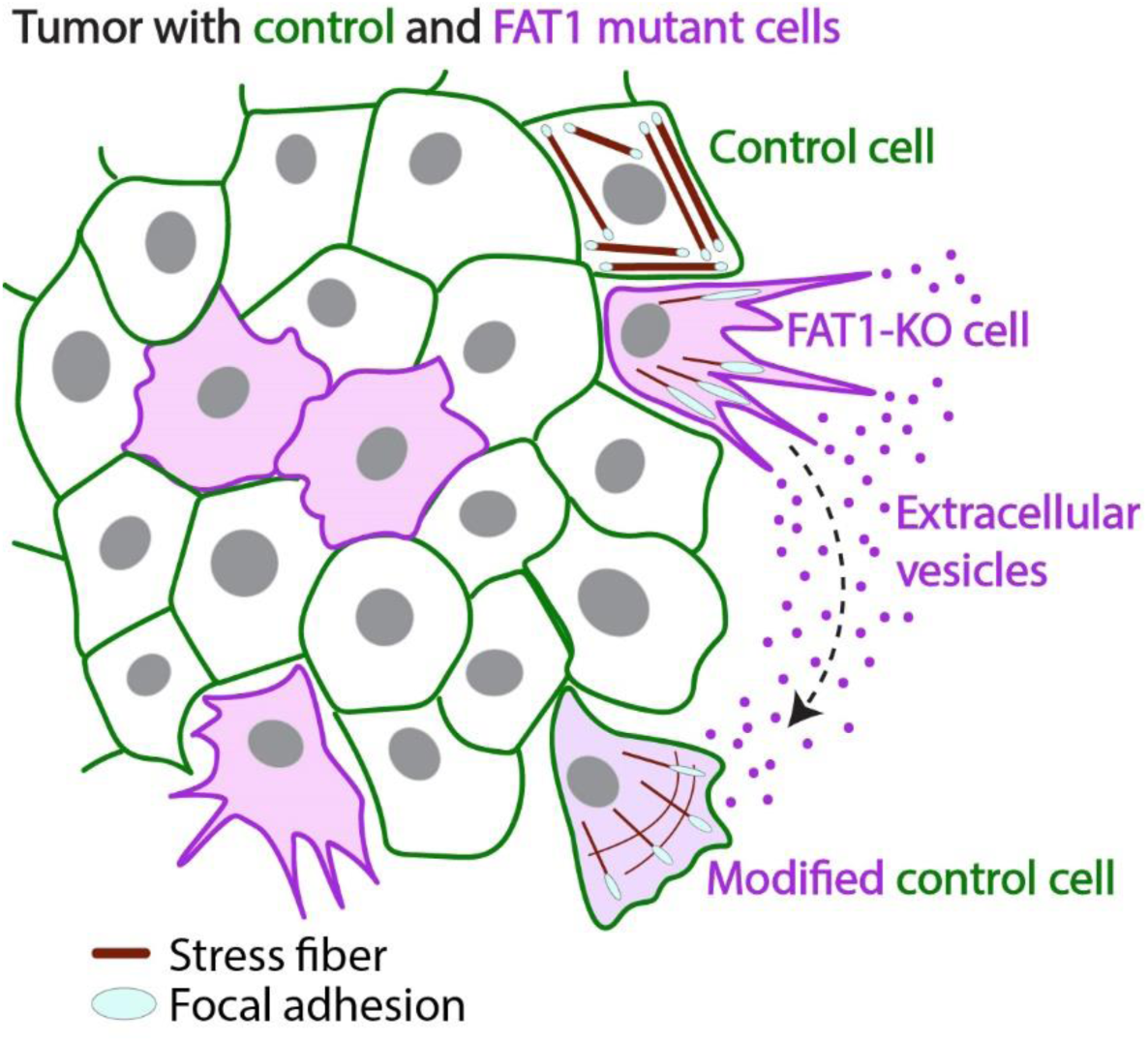
Schematic showing how FAT1 deletion stimulates the migration of neighboring non mutant cells. FAT1 KO cells secrete extracellular vesicles that carry factors that promote migration of non-mutant control cells, possibly through relaxation of traction forces in the recipient cells.

## Introduction

Cancer metastasis is a complex and dynamic process that involves uncontrolled cell growth, evasion from immune cells, and the acquisition of motile properties that allow for tumor dissemination across the body (Hanahan and Weinberg, 2011). Numerous cancer cells undergo a gradual, dynamic and reversible transition between epithelial and mesenchymal states, which is in fact a normal developmental process that is reactivated during tumor progression (Cheung and Ewald, 2016; Nieto et al., 2016; Thiery et al., 2009; Tsai et al., 2012). During epithelial-to-mesenchymal transition (EMT), epithelial cells lose cell-cell contacts and junctional integrity through the loss of adhesion proteins such as E-cadherin (E-Cad), occludins and claudins, and express transcription factors including *ZEB1, SNAIL* and *TWIST* that simultaneously allow for activation of mesenchymal markers including vimentin, N-cadherin and matrix metalloproteases (Yang et al., 2020). In addition, epithelial cells change from a cuboidal to an elongated shape, as well as turn their apico-basal polarity into a front-rear polarity that facilitates cell migration and invasion through degradation of the extracellular matrix (Bakir et al., 2020; Brabletz et al., 2021). This process of cell scattering, polarity remodeling and acquisition of motile phenotypes is mediated through complex, and yet only partially characterized crosstalk mechanisms involving the cytoskeleton networks and cell adhesion proteins.

Interestingly, in healthy adult tissues, neither epithelial nor mesenchymal cells are invasive. Therefore, how the transition between these two states can induce cell invasion and metastasis is a key, yet unexplained consequence of EMT in tumors. It has become increasingly evident that a subpopulation of tumoral cells express both epithelial and mesenchymal markers (**Figure 1A**). These states are not transient but rather represent a stable “hybrid” EMT state (Kröger et al., 2019; Pastushenko et al., 2018), which is associated with increased invasiveness and poor survival outcomes (Brown et al., 2022; Jolly et al., 2016; Kröger et al., 2019; Simeonov et al., 2021). Thus, it is straightforward to hypothesize that these hybrid EMT cells have specific migratory properties and are responsible for the scattering and invasion of tumor cells (Jolly et al., 2018, 2015). Identification of the differentially expressed genes in hybrid EMT revealed that cells have altered expression levels of several cytoskeletal and adhesion proteins (Bocci et al., 2021; Cook and Vanderhyden, 2020; Deshmukh et al., 2021; Pastushenko et al., 2021, 2018), that might play a key role in the acquisition of their invasive properties. However, the cellular mechanisms conferring highly motile capacities to hybrid EMT cells are unknown.

**Figure 1:**
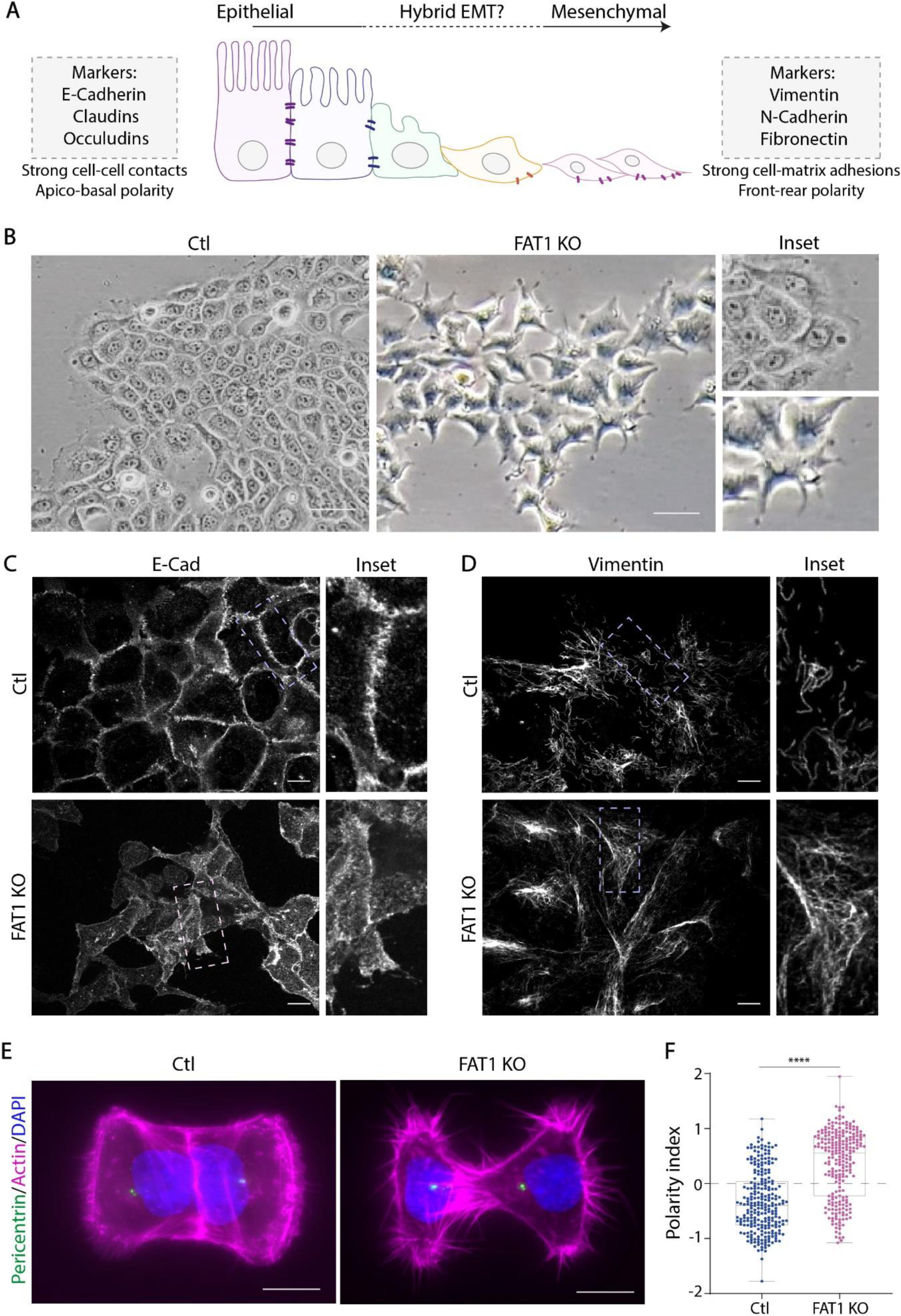
FAT1 knock-out (KO) cells possess hybrid EMT markers and the lack of polarity. (A) Epithelial-to-Mesenchymal transition can be divided in three states: Epithelial, Hybrid and Mesenchymal. Cells in epithelial states express epithelial markers as E-cadherin, stable microtubules, and apical-basal polarity. The mesenchymal state is characterized by the expression of markers such as Vimentin and N-cadherin. In addition, mesenchymal cells are characterized by their migratory behavior and loss of polarity. Interestingly, there is an emergent hybrid state where cells acquire mesenchymal characteristics and markers, and yet, maintain their epithelial markers. (B) Brightfield images of wild-type cells and FAT1 KO cells showing morphological differences. Images were acquired by brightfield microscopy using a 10× objective. Scale bar: 100 μm. (C) Confocal images of cells stained with E-Cadherin, showing a maximum intensity projection of 10 slices, spaced 1 micron, and zoomed images are shown on the right. Scale bar: 10 μm. (D) Confocal images of cells stained with Vimentin, showing at a maximum intensity projection of 10 slices, spaced 1 micron, and zoom images are shown on the right. Scale bar: 10 μm. (E) Confocal images of doublet of control or KO cells on H shaped micropatterns, size 1100 mm², stained for F-actin, pericentrin and DAPI, at a maximum intensity projection of 8 slices, spaced 1 micron. Scale bar: 20 μm. (F) Graph represents polarity index measurements of centrosome positioning in control and KO cells. When 0.5<PI<2, centrosome is close to the cell-cell junction; when -0.5>PI>-2, centrosome is at the opposite side of cell-cell junction. Horizontal lines indicate the median value for each cell type. Represented data shown are from three independent experiments, for which n was between 50 and 80 cells for each cell type. ****p < 0.0001 by Mann-Whitney T-Test.

The study of hybrid EMT cancerous cells is limited by the necessity to isolate, sort, and maintain in culture primary cells extracted from invasive tumors (Pastushenko et al., 2018). A recent study showed that the loss of FAT1, a proto-cadherin and component of cell-cell junctions, can induce a stable hybrid phenotype in mouse models (Pastushenko et al., 2021). On the other hand, contradictory studies have shown that FAT1 knockdown or deletion might be linked to reduction in EMT (X. Hu et al., 2017; Srivastava et al., 2018). FAT1 is one of the most commonly mutated genes in human cancers, particularly in squamous cell carcinomas and head and neck cancers (Dotto and Rustgi, 2016; Lawrence et al., 2014; Lin et al., 2014). In addition, FAT1 deletion is strongly associated with increased metastasis (Lan et al., 2022; Pastushenko et al., 2021; Wang et al., 2019), possibly through its effects on controlling cell growth and cell cycle genes including YAP, Wnt and CDKs (Lan et al., 2022), as well as changes in actin dynamics and cell adhesion through Src phosphorylation and VASP recruitment to actin complex and to the leading edge of cells (Moeller et al., 2004; Pastushenko et al., 2021; Tanoue and Takeichi, 2004). Therefore, given the above findings, the deletion of FAT1 offers an opportunity to study the motility of hybrid EMT cells and understand their role in tumor dissemination.

## Results

### FAT1 deletion induces hybrid EMT in squamous carcinoma cells

We investigated the cytoskeletal architecture and motility of hybrid EMT cells by deleting the FAT1 gene in a squamous cell carcinoma cell line, the A388. We utilized CRISPR-Cas9 based editing technologies to delete FAT1 and obtained two distinct clones that we characterized in parallel throughout this study (Supplementary Fig. 1A). In culture, the control wild-type cancer cells formed dense colonies that were highly cohesive, whereas the FAT1 KO cells formed sparse clusters with individual cells being star-shaped and more loosely connected to their neighbors (**Figure 1B**). As expected from a previous study (Pastushenko et al., 2021), FAT1 KO cells displayed a partial decrease of E-cadherin expression (**Figure 1C**), and an upregulated level of vimentin, a classical mesenchymal marker, which formed an extended network of filaments (**Figure 1D**, Supplementary Figure S1B). Interestingly, the analysis of cell polarity in cell doublets confined on micropatterns showed that the orientation of the nucleus-centrosome axis in control cells was determined by the positioning of nuclei toward intercellular junctions (**Figure 1E**), whereas the weakening of these junctions in FAT1 KO led to nucleus repositioning toward cell adhesions and away from the extracellular matrix, resulting in an effective reversal of cell polarity (**Figure 1F**). Polarity reversal was shown to be an early sign of the remodeling of epithelial polarity in response to TGF-beta (Burute et al., 2017) or ZEB1 expression (Margaron et al., 2019). All these results were confirmed using the second FAT1 KO clone (Supplementary Figure S1). Altogether, our results confirmed that the deletion of FAT1 induces a hybrid EMT phenotype in A388 cells.

### FAT1 deletion reduces cell migration properties

First, we investigated the invasive properties of FAT1 KO cells in 3D by studying their escape and motility from spheroids embedded in matrigel (**Figure 2A**). Control cells formed a compact and round spheroid, whereas FAT1 mutant cells formed less cohesive spheroids (**Figure 2A**). However, upon tracking cell migration, we found that although FAT1 KO cells initially detached from the spheroid, they were unable to migrate further away (**Figure 2B**, Supplementary Movie S1). This behavior might result from defective ECM proteolysis or specific hydrogel composition/stiffness. To reduce the potential contributions of multiple parameters, and further investigate this unexpected behavior, we studied cell migration in 2D using wound healing assays (**Figure 2C**). In 2D migration assays as well, we observed that the migration properties of FAT1 KO cells were impaired: they moved slower and less persistently (**Figure 2D, 2E**, Supplementary Movie S2). This latter measurement of directionality might be caused by their defective interactions with adjacent cells which is a key parameter for collective migration and wound healing. Therefore, we investigated the intrinsic migration capacities of individual cells, without any potential interference of the additional contribution of adjacent cells. To this end, we simplified our model and analyzed single cell migration in 1D using microprinted lines of fibronectin, which can recapitulate the process of cell migration along ECM fibers in 3D (Doyle et al., 2009). The results were even more striking: mutant cells could not move at all (**Figure 2F**, Supplementary Movie S3). Since cell motility can strongly depend on the level of cell adhesion and substrate stiffness (Gupton and Waterman-Storer, 2006), we varied these parameters in our model system to characterize FAT1 KO migration. In all cases, FAT1 KO cells were unable to migrate (**Figure 2G**). We also confirmed all these results using the second FAT1 KO clone (Supplementary Figure S2). Taken together, we show that hybrid EMT phenotype induced by FAT1 deletion is characterized by defective cell migration. This was surprising as FAT1 deletion was shown to promote tumor invasiveness and metastasis in mice (Pastushenko et al., 2021).

**Figure 2:**
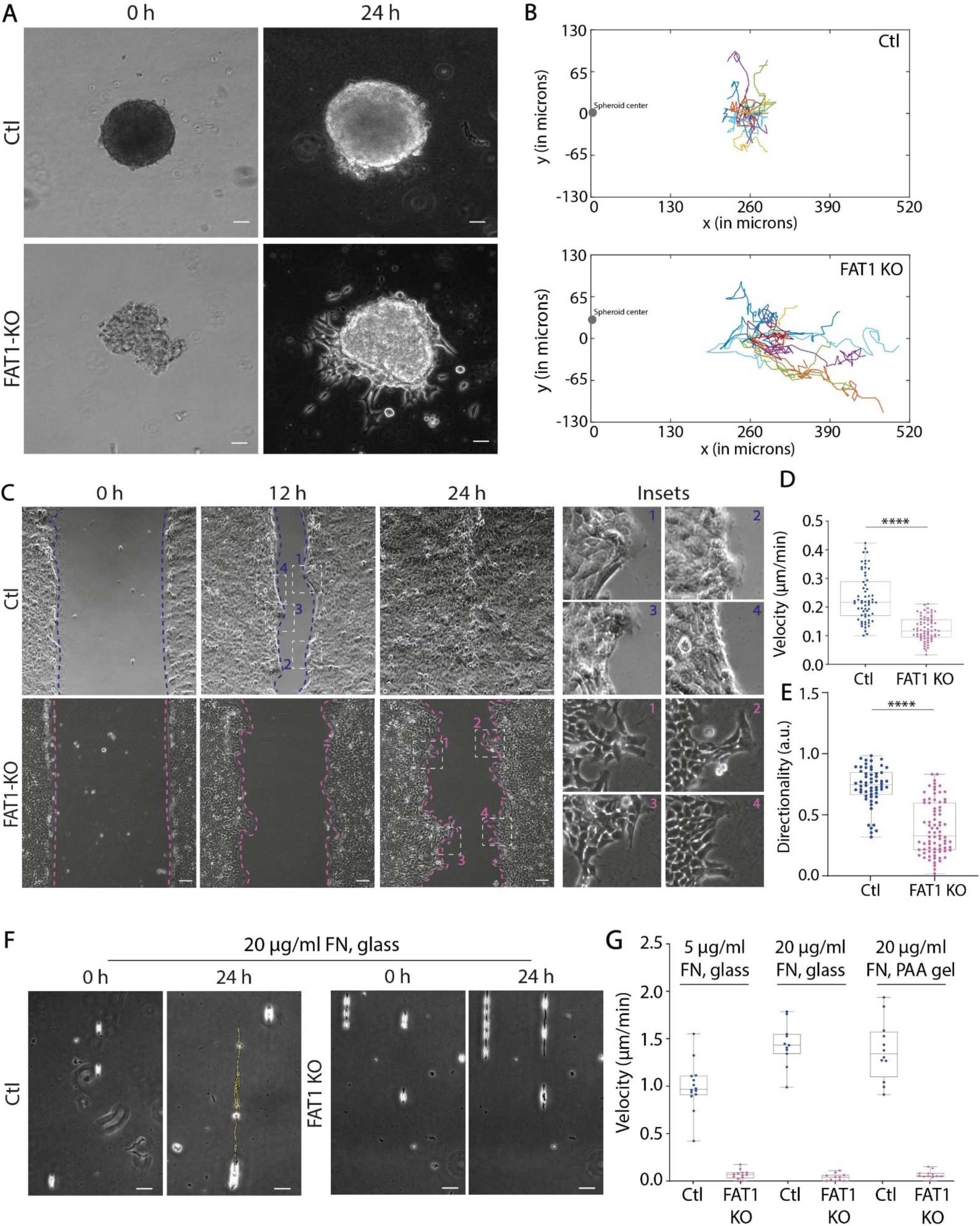
FAT1 knock-out impairs cell mobility despite hybrid EMT characteristics. (A) Transmitted light images of control and KO cells on a 3D migration assays in Matrigel at the indicating time. Images were acquired for 48 h, every 15 min. Scale bar: 100 μm. See Supplemental Videos S1. (B) Graphs represent the trajectories of escaping cells from the spheroids. (C) Transmitted light images of control and KO cells migrating at the indicated times after monolayer wounding. Images were acquired for 24 h, every 15 min. Scale bar: 100 μm. See Supplemental Videos S2. (D) Graph represents cell velocity from wound healing. Represented data shown are from three independent experiments, for which n was between 30 and 50 cells for each cell type. Graph represents cell persistence of cells. Represented data shown are from three independent experiments, for which n was between 30 and 50 cells for each cell type. Bar 100 μm. (E) Transmitted light images of single cell migrating control and KO on micropatterned lines, coated with Fibronectin at 20 μg/ml. Dotted yellow lines show the trajectories of a single cell. (F) Graph represents single cell velocity. Plotted data come from three independent experiments, for which n was between 10 and 15 cells for each cell type. See supplemental video S3. For all graphs, ****p < 0.0001 using Mann-Whitney T-Test.

### Migration properties are improved in mixed populations of control and FAT1 KO cells

Tumor masses consist of heterogenous populations of cells, where multiple cell types exhibit various mutation profiles (Januškevičienė and Petrikaitė, 2019). In this context, mechanical interaction of heterogenous populations of cells with distinct adhesion and contractile properties can lead to cell sorting (Krieg et al., 2008), formation of repulsive boundaries (Rodríguez-Franco et al., 2017) and active reorganization (Labernadie et al., 2017; Pajic-Lijakovic, 2023) which might contribute to tissue fluidization and improve collective migration (Jain et al., 2020). This led us to hypothesize that the invasiveness of tumors containing FAT1-mutant cells in mice might result from the interaction of heterogenous population of control and mutant cells. Thus, we mixed control and FAT1 KO cells in a wound healing assay to observe the collective migration of the two populations (**Figure 3A**). As the FAT1 KO cells proliferate faster, the initial density of FAT1 KO cells were half that of control cells in the migration assay to allow for similar proportions of the two cell types at the start of migration. Interestingly, the mixed population migrated faster and closed the wound earlier than a homogeneous population of control cells (**Figure 3B** and Supplementary Movie S4). Both cell types migrated faster together when mixed than individually (**Figure 3B**). The directionality of control cells was impaired, whereas that of FAT1 KO was improved in the mixed populations (**Figure 3C**).

**Figure 3:**
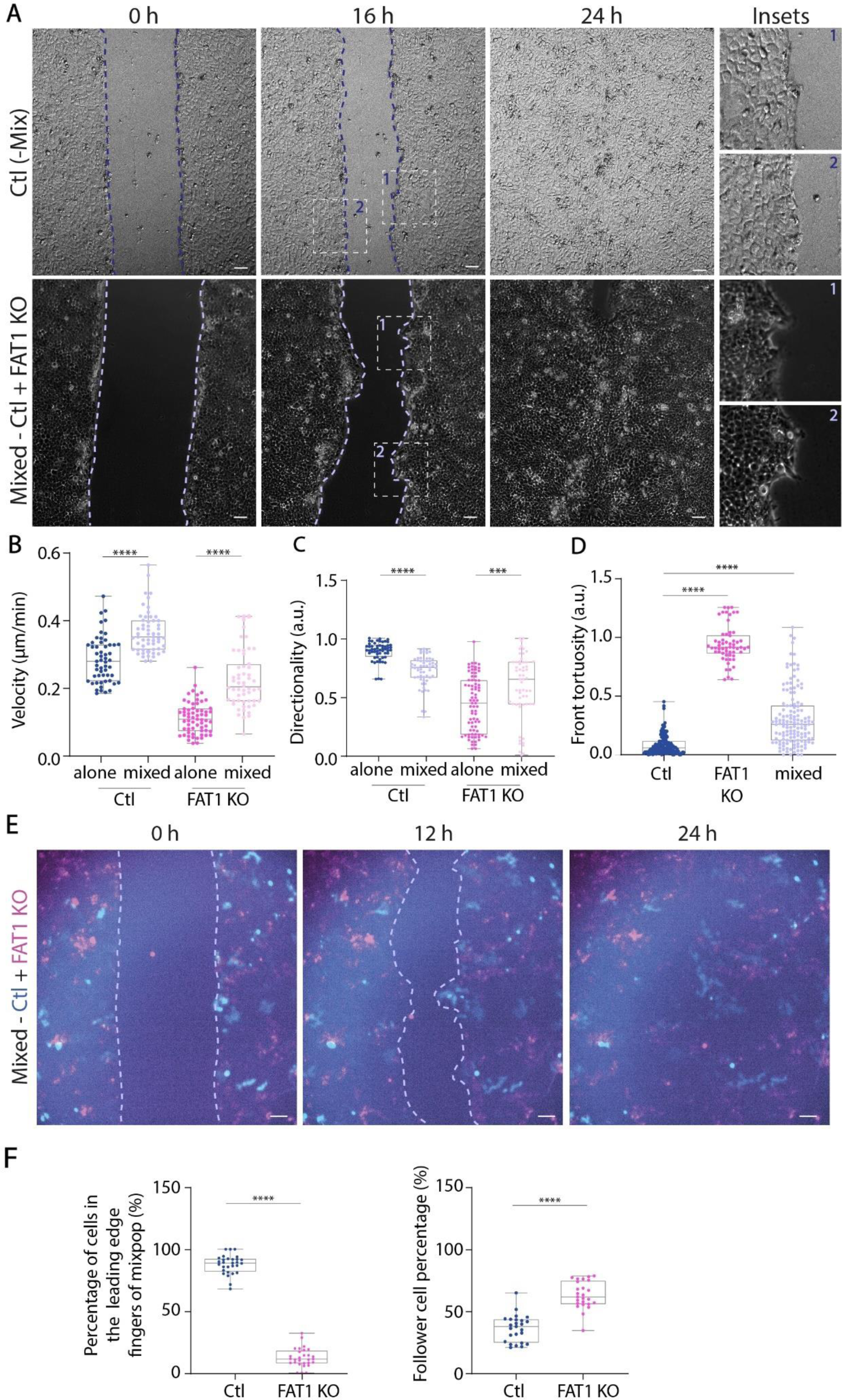
Mixing FAT1 KO and control cells improves collective migration. (A) Transmitted light images of control and mix population of control and FAT1 KO cells migrating at the indicated times after monolayer wounding. Images were acquired for 24 h, every 15 min. Scale bar: 100 μm. See Supplemental video S4. (B) Graph represents cell velocity of migrating cells from wound healing. Represented data shown are from three independent experiments, for which n was between 40 and 50 cells for each cell type. (C) Graph represents directionality of cells. Represented data shown are from three independent experiments, for which n was between 20 and 30 cells for each cell type. (D) Graph represents front tortuosity for control cells and the mix population control + FAT1 KO. Ratio between the average length of the front every 1 h and the initial length ratio represents the tortuosity of the front. Represented data shown are from three independent experiments, for which n was between 20 and 30 cells for each cell type. (E) Fluorescent images of control and mix population control + FAT1 KO at the indicated time after monolayer wounding. Control cells were stained with Orange Cytotracker and are shown in blue, FAT1 KO cells were stained with Far-Red Cytotracker and are shown in pink. Images were acquired for 24 h, every 15 min. Scale bar: 100 μm. See Supplemental video S5. (F) Graph represents cell type proportion in migration fingers. Represented data shown are from three independent experiments, for which n was between 10 and 20 fingers. For B-F, ****p < 0.0001 using Mann-Whitney T-Test.

Interestingly, the migrating front of the mixed collective was tortuous and displayed finger-like protrusions (**Figure 3E, F**, Supplementary Movie S5). Such “fingers” are mechanical instabilities of the cell front that actively contribute to promote the collective migration process (Reffay et al., 2014). We also observed such fingers in a homogeneous population of mutant cells but not in control cells (**Figure 2C, 3D**). However, surprisingly, in the case of mixed population, protrusive fingers contained mostly control cells (**Figure 3E, F**). To note, these results were confirmed in the second FAT1 KO clone (Supplementary Figure S3). Our findings suggest that the presence of mutant cells with FAT1 deletion had an impact on the mechanical properties of control cells, leading them to accelerate and form finger-like instabilities at the migrating front. Interestingly, mutant cells were not in or behind at the base of protrusive fingers, suggesting that they affect control cells from a distance rather than through direct intercellular contacts.

### FAT1 KO cells secrete factors promoting the migration of non-mutant cells

Long-range intercellular communication by secreted factors has recently emerged as a potent way for tumor cells to influence their environment (Wortzel et al., 2019). To test the potential role of secreted factors, we collected the culture medium of FAT1 KO cells after few days of cell growth and division. This “conditioned” medium contained all the factors secreted by FAT1 KO cells. We found that the addition of conditioned medium from FAT1 KO cells on control cells increased their migration velocity, but decreased their persistence and directionality, much like when FAT1 mutants were mixed with control cells (**Figure 4A-C**, Supplementary Figure S4, Supplementary Movie S6). Furthermore, the conditioned medium also induced finger-like instabilities at the front of migrating cells (**Figure 4D**). This effect could be documented by measuring either the tortuosity of the migrating front or the angle at the tip of finger-like protrusions (**Figure 4E, F**). In addition, we confirmed these findings using the second FAT1 KO clone (Supplementary Figure S4).

**Figure 4:**
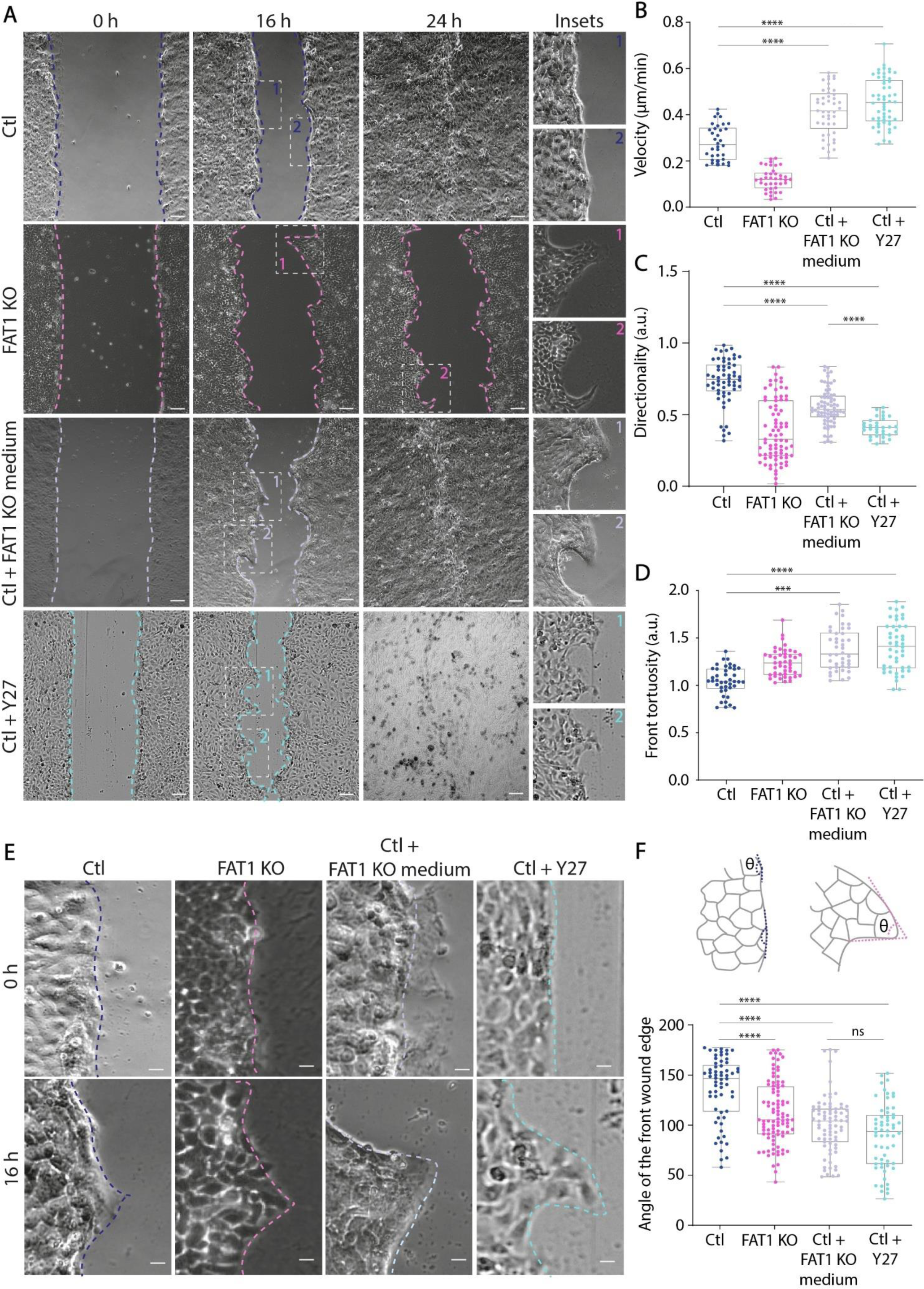
Secretion from FAT1 KO cells promotes the motility of control non-mutated cells. (A) Transmitted light images of control, FAT1 KO, control cells treated with either FAT1 KO medium or ROCK inhibitor Y-27632 (Y27) after monolayer wounding. Control cells were treated with FAT1 KO medium or Y27 one hour before wounding. Images were acquired for 24 h, every 15 min. Scale bar: 100 μm. See supplemental video S6. (B) Graph represents cell velocity of migrating cells from wound healing. Represented data shown are from three independent experiments, for which n was between 30 and 40 cells for each cell type and condition. (C) Graph represents cell directionality of cells. Represented data shown are from three independent experiments, for which n was between 30 and 40 cells for each cell type. (D) Graph represents front tortuosity for each cell type and condition. Ratio between the average length of the front every 1 h and the initial length represents the tortuosity of the front. Represented data shown are from three independent experiments, for which n was between 30 and 40 cells for each cell type. (E) Representative images of angle measurements for each cell type and condition. Angles measurements were obtained by the average of angle of the front every hour from wound healing movies. Scale bar: 10 μm. (F) Schematic of angle measurements at the wound edge. (G) Graph represents angle measurements for each cell type and condition. Represented data are from three independent experiments, for which n was between 10 and 20 migrating fingers for each cell type and condition. For all graphs, ****p < 0.0001 using Mann-Whitney T-Test.

### Pro-migratory factors are associated with extracellular vesicles

Recent studies have shown that tumor cells have a high tendency to secrete extracellular vesicles that enhance tumor growth and metastasis (Becker et al., 2016; García-Silva et al., 2021; Kalluri and McAndrews, 2023; Mathieu et al., 2019; Raposo and Stoorvogel, 2013; Wortzel et al., 2019). This prompted us to investigate which component of the secretions from FAT1 KO cells affected the migration of control cells. We collected FAT1 KO conditioned medium and sorted four fractions containing either extracellular vesicles or soluble factors using size exclusion chromatography (Royo et al., 2020) (**Figure 5A**). Conditioned medium from FAT1 KO cells had a higher concentration as well as absolute number of extracellular vesicles as compared to medium collected from control cells (**Figure 5B**). We confirmed that the fractions were composed of extracellular vesicles by western blotting for markers CD9, CD63, Synt-1 and Alix (**Figure 5C**). Fractions containing extracellular vesicles or soluble factors from FAT1 KO medium were then tested on control cells during wound healing (**Figure 5D**). We found that the addition of the fraction containing extracellular vesicles to control cells accelerated their migration velocity, decreased motion persistence, and induced finger-like instabilities at the migrating front (**Figure 5E, F, G**, Supplementary Movie S7). By contrast, the addition of the fraction containing soluble factors had no detectable effect (**Figure 5E, F, G**, Supplementary Movie S8). These results suggest that the signals in FAT1 KO conditioned medium that promoted the migration of control cells were not soluble factors, but were in fact, extracellular vesicles.

**Figure 5:**
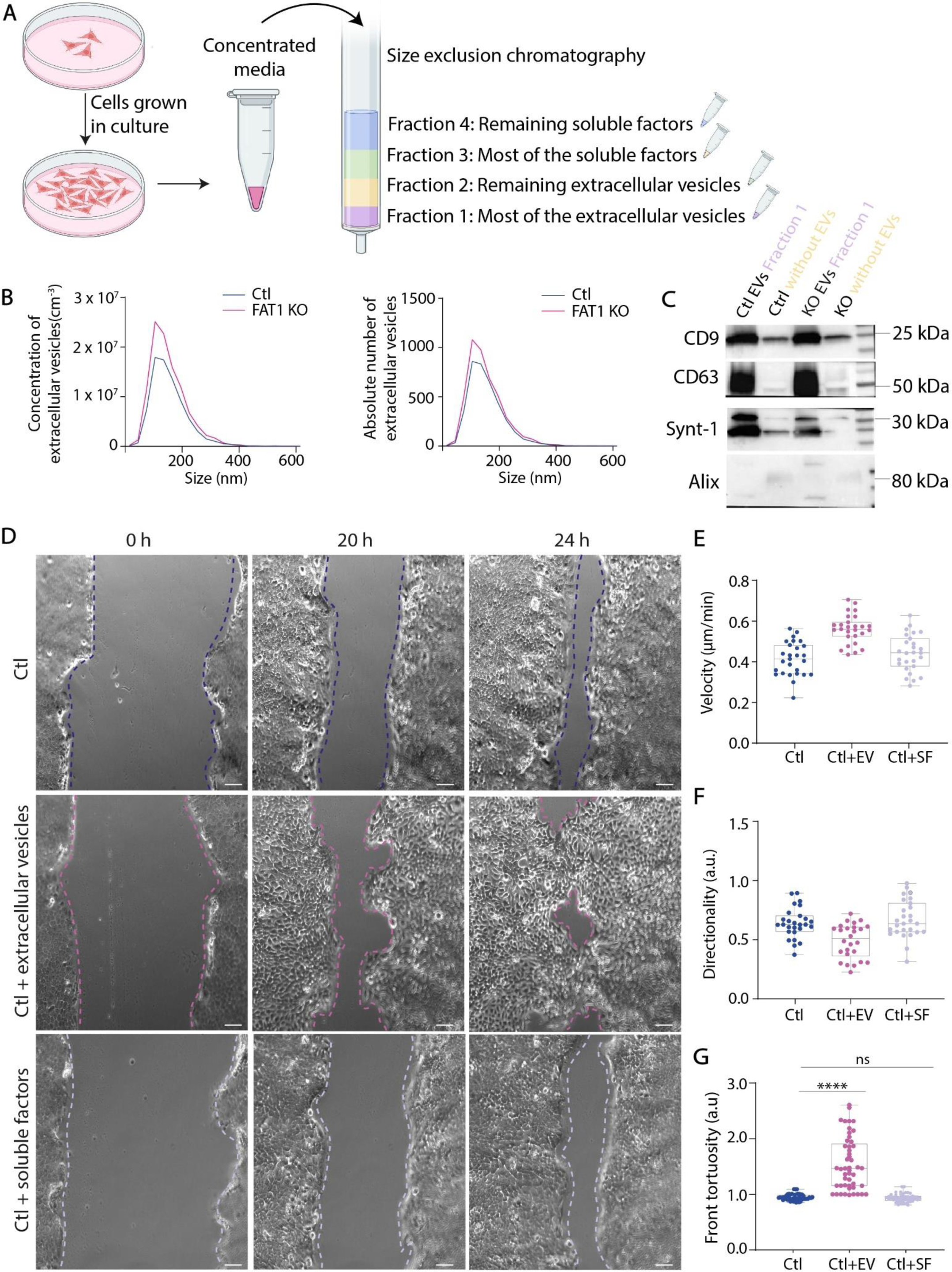
Secretion from FAT1 KO cells reduce contractility of control cells. (A) Confocal images of cells stained with F-actin staining in cells, showing a maximum intensity projection of 11 slices, spaced 1 micron, zoom images on the right for the control treated with FAT1 KO medium. Scale bar: 10 μm. (B) Graph represents proportion of control cells periphery occupied by actin spikes. Represented data shown are from three independent experiments, for which n was between 5 and 20 images for each cell type and conditions. (C) Confocal images of cells stained with pMLC, showing a maximum intensity projection of 11 slices, spaced 1 micron. Scale bar: 10 μm. (D) Graph represents pMLC intensity in each condition. Represented data shown are from three independent experiments, for which n was between 30 and 40 cells for each cell type and condition. (E) Scheme represents TFM principle, using 550 nm beads to measure the mechanical energy of a single cell on a rectangle-shaped micropattern. (F) Stress-field maps of control or FAT1 KO cells on rectangle-shaped micropatterned PAA gels of 20 kPa rigidity. (G) Graph represents mechanical energy released by each cell type. Represented data shown are from three independent experiments, for which n was between 30 and 40 cells for each cell type and condition. (H) Confocal images of cells stained with paxillin, showing a maximum intensity projection of 8 slices, spaced 1 micron. Scale bar: 10 μm. For all graphs, ****p < 0.0001 using Mann-Whitney T-Test.

### Pro-migratory factors reduce cell contractility

Finally, we investigated the mechanism by which FAT1 deletion and the pro-migratory secreted factors affect the mechanical properties of control cells, destabilize their migrating front, and increase their migration velocity. First, we characterized the organization of the actomyosin network in our system. We found that control cells displayed numerous contractile fibers: stress fibers that occupy the entire ventral cell surface, thick transverse arcs connecting radial fibers, and conspicuous peripheral cables at the wound edge suggestive of large intercellular tensional forces (Chen et al., 2019) (**Figure 6A**). On the one hand, such intercellular tensional forces can promote wound closure (Chen et al., 2019; Tambe et al., 2011) but on the other hand, high traction forces on ECM promote the growth of focal adhesions, which reduces their necessary turnover and detachment, thus, stalling cell migration (Hennig et al., 2020). FAT1 KO cells exhibited different actomyosin structures: contractile fibers were much less prominent; stress fibers had almost disappeared and were replaced by numerous filopodia at the cell surface (**Figure 6A**). The presence of these filopodia was consistent with the overactivation of Src and FAK in FAT1 KO cells (Jacquemet et al., 2015; Pastushenko et al., 2021). Consistent with the observations of these various actin-based structures, we found that phospho-myosin light chain (pMLC) intensely concentrated along ventral fibers in control cells (S. Hu et al., 2017), and localized at the base of filopodia at the dorsal cell surface although with lower concentration (Alieva et al., 2019) (**Figure 6B, C**). All these features were suggestive of the actin network being more involved in filopodia assembly and dynamics at cell dorsal surface, rather than the production of contractile forces along the ventral surface in FAT1 KO cells. Interestingly, control cells treated with FAT1 KO conditioned medium showed a mixed phenotype: stress fibers were still present but thinner, and numerous filopodia were visible at the cell surface (**Figure 6A, B**).

**Figure 6:**
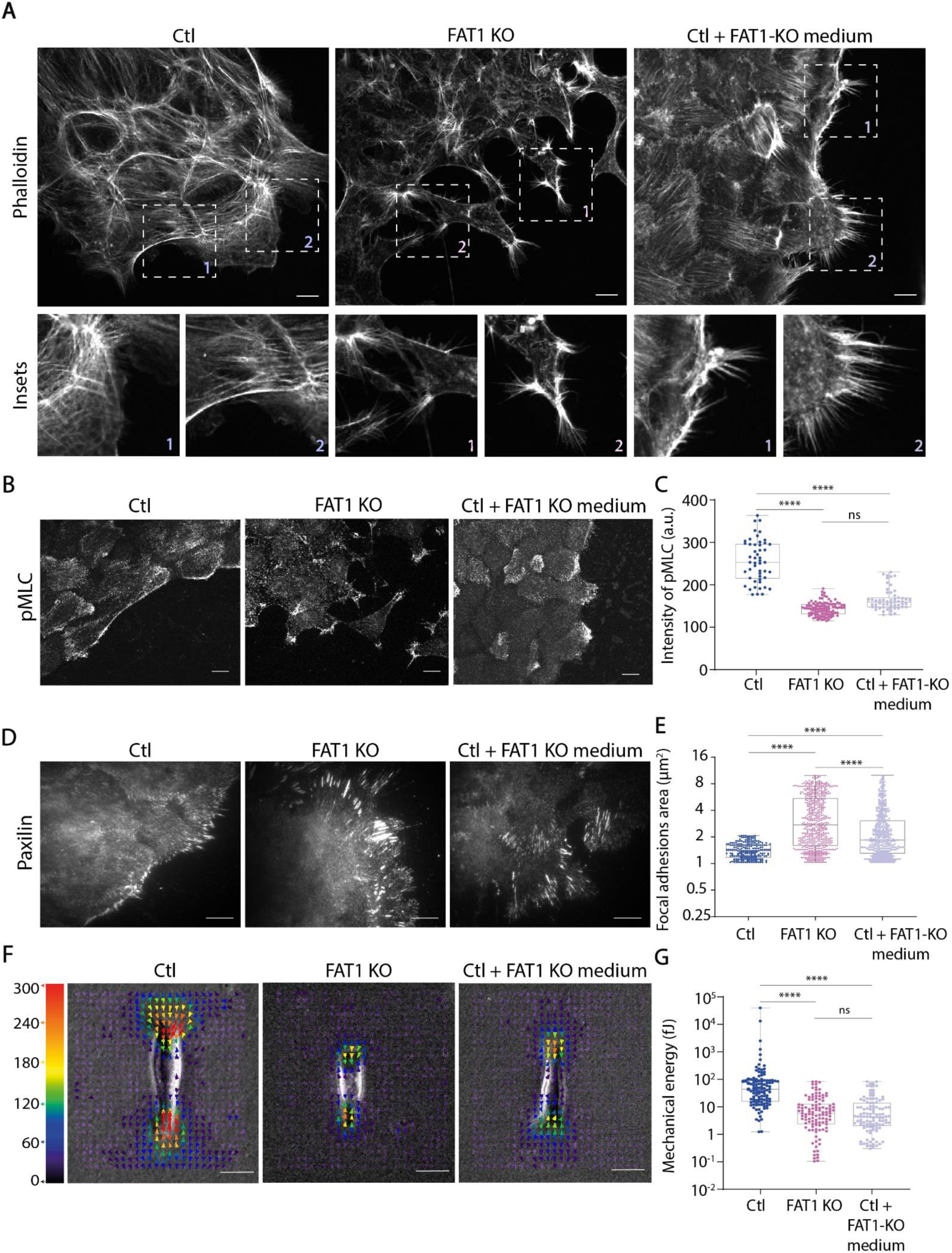
Secretion from FAT1 KO cells could be extracellular vesicles. (A) Schematic represents SEC technique to extract extracellular vesicles (EVs) and soluble factors from cell medium. (B) Graphs represent the concentration of EVs and absolute number extracted from control and FAT1 KO medium. (C) Transmitted light images of control cells migrating at the indicated times after monolayer wounding. Control cells were treated with different extracted fractions (EVs and soluble factors) from FAT1 KO medium one hour before wounding. Images were acquired by time-lapse microscopy for 24 h, time point 15 min. Scale bar: 100 μm. See supplemental video S7. (D) Graph represents cell velocity of migrating cells from wound healing. Represented data shown are from three independent experiments, for which n was between 30 and 40 cells for each cell type and condition. (E) Graph represents cell directionality of cells. Represented data shown are from three independent experiments, for which n was between 30 and 40 cells for each cell type. (F) Graph represents front tortuosity of cells. Represented data shown are from three independent experiments, for which n was 20 images for each cell type. For all graphs, ****p < 0.0001 using Mann-Whitney T-Test.

Surprisingly, the observation of paxillin staining to reveal focal adhesions, the size of which is supposed to be positively correlated to the magnitude of traction forces produced by cells on their environment (Kollimada et al., 2021), revealed a different picture: focal adhesions were rare and quite small in control cells resembling an epithelial colony edge, whereas FAT1 KO cells exhibited numerous and extremely long focal adhesions (**Figure 6D, E**). In addition, control cells treated with FAT1 KO medium exhibited bigger focal adhesions than the non-modified cells (**Figure 6E**). To solve this apparent contradiction, we directly measured the magnitude of traction forces produced by cells on their environment as they migrate. To this end, we combined cell micropatterning on deformable substrates to normalize cell shape and traction force microscopy (Kollimada et al., 2021; Kurzawa et al., 2017). We show that the traction forces generated on the extracellular matrix were almost an order of magnitude lower in FAT1 KO cells. Furthermore, we found that the addition of FAT1 KO conditioned medium on control cells was sufficient to reduce their contractility to the level of FAT1 KO cells (**Figure 6F**). All these data were confirmed in the second FAT1 KO clone (Supplementary Figure S5). These results showed that FAT1 deletion dramatically reduced the production of traction forces on cell environment and that the factors secreted by FAT1 KO cells contained signals capable of reducing the contractility of recipient cells. This conclusion was further supported by the observation that the downregulation of control cell contractility by an inhibitor of Rho kinase (Y-27632) had similar effect as the addition of FAT1 KO conditioned medium, i.e., increase of cell velocity in wound healing (**Figure 4B**), reduction of motion directionality (**Figure 4C**) and induction of destabilizing fingers along the migrating front (**Figure 4D, F**). Altogether, our findings suggest that FAT1 KO cells influence non-mutated squamous carcinoma cells that are quite contractile, by the secretion of factors that reduce their contractility.

## Discussion

In this study, we discovered that the hybrid EMT cells induced by the deletion of FAT1 protocadherin in carcinoma cells were poorly capable of migrating in 1, 2 or 3D. These observations were extremely surprising considering the direct role of FAT1 deletion (Pastushenko et al., 2021) and the more general role of hybrid EMT cells (Kröger et al., 2019) in the induction of tumoral invasiveness and metastasis. However, an additional series of experiments led us to uncover a possible explanation for this discrepancy. We found that FAT1 KO cells impart invasive properties to the surrounding non-mutated cells by secreting pro-migratory signals. We further identified that the pro-migratory signals were associated with extracellular vesicles, and are capable of reducing the high contractility of surrounding cancer cells. This showed that while FAT1 KO cells may not be capable of invading themselves, they can secrete factors that force other cells to become invasive. Our observations add to the growing list of non-cell autonomous functions of genes that play a crucial role in altering signaling pathways governing tumor growth, proliferation and metastasis (Corso and Giordano, 2013; Lujambio et al., 2013; Tissot et al., 2017).

How does a reduction of contractility promote the migration of cancer cells? Mesenchymal-like migration of adhesive cells requires a subtle balance of adhesion strength and contractile forces. However, these are not linear processes, but in fact, function optimally at intermediate levels of adhesion and contraction (Gardel et al., 2010, 2008; Gupton and Waterman-Storer, 2006; Schwarz and Gardel, 2012; Seetharaman and Etienne-Manneville, 2020; Wisniewski et al., 2020). A minimal level of adhesion is necessary for cells to attach and translocate. In parallel, contractility needs to be sufficient to deform cell body. However, when contractility is too high, contractile forces promote the enlargement of cell adhesions that can no longer be detached from the matrix, and thus, prevent cell displacement (Leal-Egaña et al., 2017). The migration of squamous carcinoma cells in our study was probably limited by the elevated magnitude of traction forces that we recorded. By contrast, traction forces were extremely low in FAT1 KO cells. It is possible that the vesicles secreted by FAT1-mutant cells likely included specific signals responsible for lowering the contraction of the cortical actin network. As these vesicles interact and fuse with non-mutated cells, they possibly transfer those signals and thereby, reduce the contractility of non-mutated cells and promoted their migration.

Extra-cellular vesicles from FAT1 KO cells reduced contractility of control cells to the level of mutant cells and promoted their migration. However, this seems counter-intuitive because despite having a similar level of traction forces, the FAT1 mutant cells were unable to migrate. The explanation might reside in the fact that changes in the number and size of their focal adhesions might alter focal adhesion turnover in FAT1 deleted cells. Mesenchymal migration is initiated and limited by the detachment of cell adhesions at the rear (Hennig et al., 2020). The adhesion we observed in mutant cells does not seem to be compatible with the required detachment and the corresponding turnover of focal adhesions that is necessary for cell migration. In particular, such large focal adhesions are highly reminiscent of adhesions generating low traction forces we measured, preferentially expressing β_1_ over α_V_ (Schiller et al., 2013).

Interestingly, FAT1 KO displayed an impressive amount of filopodia, which likely contributed to production of extracellular vesicles. The high level of Src phosphorylation in FAT1 KO (Pastushenko et al., 2021) is consistent with the reorganization of the actomyosin network into the assembly of filopodia (He et al., 2015). In addition, the low level of traction forces in these cells might also be related to the relocalization of myosin II from stress fibers to the base of filopodia (Alieva et al., 2019). Filopodia are thin dynamic protrusions which also act as platforms for the shedding of extracellular vesicles (Rilla, 2021). Their thin shape can easily undergo pearling instabilities and generate numerous vesicles. They can also be actively produced by the pinching of filopodia tip (Nishimura et al., 2021). Filopodia fragmentation has been documented in several cancer cells (Arasu et al., 2017; Obregon et al., 2006; Rilla, 2021; Shibue et al., 2012). Such filopodia-derived vesicles and other small extracellular vesicles have also been shown to promote the migration of recipient cells (Sneider et al., 2023), notably by transferring active Rac (Nishimura et al., 2021). Extracellular vesicles from cancer cells have also been shown to prepare the pre-metastatic niche to disseminate tumors to surrounding tissues (García-Silva et al., 2021; Tkach and Théry, 2016). These observations along with our results suggest that the remodeling and weakening of the actin network in response to specific oncogenes might participate in tumor propagation by generating extracellular vesicles carrying actomyosin relaxing signals to promote the migration of surrounding non-mutated cancer cells.

## Materials and Methods

### Cell culture

Squamous carcinoma cells (SCC) from skin were a gift by Prof. Bernard Weissman, UNC Lineberger Comprehensive Cancer Center, North Carolina and Prof. Cédric Blainpain (Université Libre de Bruxelles). Cells were grown in 1 g/l glucose DMEM (Gibco) supplemented with 10% FBS (VWR), 1% antibiotics-antifungal solution (Fisher Scientific) at 37°C and 5% CO_2_.

### Generation of FAT1 KO cells using CRISPR/Cas9 editing

FAT1 knock out cells were generated using a pool of 3 different plasmids (Santa Cruz Biotechnologies) containing GFP and the following guide RNA (gRNA) sequences: 5’-GTCTCATCACAACTACGTCA-3’; 5’-GTGGCGTCTTCTACATGTCG-3’; 5’-GTGATAGAGCGGCTCCCGTC-3’.

WT cells were transfected with the sgRNA-containing plasmids by electroporation with a Nucleofector device (Lonza; in Mirus transfection solution). Cells were plated in culture dishes and maintained for three days. GFP-positive cells were subsequently collected using fluorescent activated cell sorting (FACS) and grown until confluency. FAT1 knock out was determined using western blot analysis and DNA sequencing. DNA extraction was made using Phenol-Chloroform method and PCR was performed using the following forward and reverse primers respectively: 5’-CTTGACCGTGAAACAACAGAC-3’ and 5’-CCACTCCAGTTTTCGGTTCG-3’. The PCR product was run on an SDS gel and purified using the Agarose Gel DNA extraction kit from Merck (11696505001).

### Western blotting

Cell lysates were obtained with SDS supplemented with Laemmli Buffer 2x (Sigma). Samples were boiled for 10 min at 94°C before loading on polyacrylamide gels (4-15% Bis-Tris Gel, Bio-Rad). Proteins were separated by gel electrophoreses at 80 V and transferred at 90 V for 2 h on nitrocellulose membranes (Precast Protein Gels, Cat. No. 4561084 and 0.2 µm nitrocellulose membrane, Cat. No. 1620112). Membranes were blocked with a solution of 5% BSA in Tris-buffered saline, 0.1% Tween 20 detergent (TBS-T). The membranes were incubated for 1 h at room temperature with primary antibody, and overnight at 4°C with horseradish peroxidase (HRP)-conjugated secondary antibody. Both primary and secondary antibody solutions were made in a solution of 5% BSA in TBS-T. Protein bands were revealed with ECL chemoluminescent substrate (Biorad) and signals were recorded using a ChemiDoc MP Imaging System (Biorad). The following primary antibodies were used for western blot analysis: anti-GAPDH (1:10000, rabbit polyclonal, sc-25778, Santa Cruz), anti-FAT1 (1:500, MABC612, mouse monoclonal, Merck). Secondary antibodies used were HRP donkey anti-rabbit and goat anti-mouse (both 1:1000, Invitrogen).

### Micropatterning

Coverslips were cleaned with acetone and isopropanol before plasma-cleaned for 5 min and incubated with 0.1 mg/ml poly-lysine/polyethylene glycol (PLL-PEG) diluted in 10 mM HEPES for 30 min at room temperature. Excess PLL-PEG was washed off coverslips using distilled water, and the coverslips were then dried and stored at 4 °C overnight before printing. Micropatterns were printed on prepared PLL-PEG coverslips for 5 min with designed chrome masks and coated with 50:50 (v/v) fibronectin/fluorescent fibrinogen mixture (20 µg/ml each) and collagen 5 µg/ml diluted in fresh 100 mM NaHCO_3_ (pH 8.3) for 30 min at room temperature. Micropatterned coverslips were washed thrice in NaHCO_3_, and incubated for 30 min with culture medium before use. Plated cells (approximately 30000 cells/ml for control cells and 15000 cells/ml for FAT1 KO cells) were allowed to adhere for 24 h before fixation. In Fig. 1F, H shaped patterns have an area of 1100 µm², in Fig.5D rectangle patterns have an area of 650 µm^2^.

### Traction force microscopy

Coverslips for traction force experiments were plasma-cleaned for 3 min and silanized for 10 min using a solution of 1% (v/v) 3-(trimethoxysilyl)propyl methacrylate and 1% (v/v) acetic acid in ethanol. The coverslips were then washed twice with absolute ethanol and dried.

PEG covalent coated beads were prepared the day of the experiment as follows: 20 µl fluorescent microbeads (555 nm) were resuspended in 10 mM MES, pH 5.5 and sonicated 30 s. Next, 500 µl of PLL-PEG (diluted in 0.1 mg/ml in 10 mM HEPES, pH 8.5) was added to the resuspended beads and the mixture was vortexed and sonicated for 1 min. A solution of EDC (1-(3-Dimethylaminopropyl)-3-Ethylcarbodiimide, 8mg/ml, Sigma, 8.00907) and NHS (N-Hydroxysuccinimide, 16 mg/ml, Sigma, 130672) was added to the fluorescent beads solution and incubated for 1 h at room temperature. The solution was centrifuged twice at 13000 rpm for 15 min. The pellet was then resuspended in 10 mM HEPES pH 7.4 to a final volume of 40 µl of beads solution. After printing and coating micropatterned coverslips, polyacrylamide gel mixtures were prepared as follows: 20 kPa – 100 µl 40% acrylamide (Bio-Rad), 66 µl 2% bisacrylamide (Bio-Rad), 334 µl water for a final concentration of 12.5% acrylamide and 3.75% bisacrylamide in water. Beads were then mixed with 145 µl of the gel mixture. After adding 1 µl APS and 1 µl TEMED, the gel mixture was added as a 25 µl drop on a silanized 20 x 20 mm coverslip. The protein-coated micropatterned coverslip was then gently placed on the gel mixture. The solution was allowed to polymerize for 30 min at room temperature. After detaching the micropatterned glass from the polymerized gel, 6 x 10^4^ cells/ml were plated on the gels and allowed to adhere and spread for 24 h before TFM experiments. Images of single micropatterned cells were acquired before and after treatment with trypsin. Acquisitions were performed with a Spinning Disk, with a 20× 0.8 NA air objective equipped with a pco.edge sCMOS camera (Orca-Flash 4.0) and Micromanager software. Cells were maintained at 5% CO_2_ and 37°C in SCC medium during acquisition. For the conditioned medium experiments, WT cells were incubated 1 h in FAT1 KO medium before experiment.

### Spheroid assays

Spheroids were produced by the hanging drop method (Foty, 2011). Cells (approx. 1000) were allowed to aggregate in a droplet of medium on the lid of a petri dish. The lid with droplets were flipped over to cover the petri dish containing PBS. The droplets containing cells are allowed to hang by gravity overnight at 37°C. The next day, each droplet containing spheroids were picked up gently using a pipette and mixed with 75% matrigel (Cat. No. 356234, Corning). The spheroids were then plated on glass-bottom wells coated with a solution of matrigel in medium (1:3). The matrigel containing the spheroid was allowed to polymerize for 30 min at 37°C, and 1 ml medium was then added in each well. The spheroids were imaged for 48 h using a video Olympus microscope. Images were acquired every 30 min with an air objective 10× 0.45 NA equipped with a pco.edge sCMOS camera (Orca-Flash 4.0) and Micromanager software.

### Migration assay

Cells were allowed to grow on defined supports (plates or coverslips) to confluence, and fresh medium was added the day of the experiment. The cell monolayer was then scratched with a p200 pipette tip to induce migration. For wound healing assays with conditioned medium, control cells were plated on plates and allowed to grow confluence. Conditioned medium was obtained after 48 h from a culture of FAT1 KO cells. Control medium was removed, and control cells were washed once with PBS before adding 1 ml conditioned medium, 1 h before inducing a wound. For mix population wound healing assays, control cells were mixed with FAT1 KO cells in a 2:1 ratio. Image acquisition was launched about 30 min after wounding or 1 hour after wounding for Conditioned medium experiments. Movies were acquired with a Video Olympus microscope equipped with a thermostatic humid chamber with 5% CO_2_ and 37°C. Images were acquired every 15 min for 24 h by an air objective 10× 0.45 NA equipped with a pco.edge sCMOS camera (Orca-Flash 4.0) and Micromanager software.

### Immunofluorescence

Migrating cells (for 8 h), sparse or a monolayer of cells, or cell spheroids were either fixed with 4% PFA, 0.25% Triton, 0.2% Glutaraldehyde 25 µM in cytoskeleton buffer for 10 min at 37°C, washed thrice with PBS, and quenched with Sodium Borohydride for 10 min at 4°C, or fixed with ice-cold methanol for 3 min at -20°C. Cells were washed thrice in PBS and stored at 4°C. Coverslips were blocked for 30 min with 5% BSA in PBS. Primary and secondary antibody were incubated 1 h at room temperature in 5% BSA. Coverslips were incubated 10 min with DAPI. Coverslips were mounted with Mowiol mounting medium (Cat. No. 81381, Sigma). Fluorescence images were acquired with a Spinning Disk microscope equipped with 40× 1.25 NA or 63× 1.4 NA objectives and recorded on a SCMOS camera (BSI) with Metamorph software.

### Antibodies, Celltrackers and drugs

Primary Antibodies used in this study are: Pericentrin (1:1000, ab4448, rabbit polyclonal, Abcam), anti-α-tubulin (1:400, ab18251, rabbit polyclonal, Abcam), anti-E-cadherin (1:200, 610181, mouse monoclonal, BD Biosciences), anti-FAT1 (1:200, HPA001869, rabbit polyclonal, Sigma Aldrich), anti-phospho myosin light chain pMLC (Ser19, 1:200, 3671, polyclonal rabbit, Cell Signaling Technology), Vimentin (D21H3, 1:400, 5741S, rabbit monoclonal, Cell Signaling Technology), Alexa Fluor 555 Phalloidin (1:400, A34055, Invitrogen), Alexa Fluor 647 Phalloidin (1:400, A22287, Invitrogen).

Secondary antibodies (1:400) used were: Alexa Fluor 488 Goat anti-Rabbit (A11008), Alexa Fluor 568 Goat anti-Rabbit (A11036), Alexa Fluor 488 Donkey anti-Mouse (A21202), Alexa Fluor 555 Donkey anti-Mouse (A31570), Alexa Fluor 647 Donkey anti-Mouse (A31571) from Invitrogen.

Cells were incubated with Celltrackers Green CMFDA Dye (C2925, Fisher) and far red (C34565, Fisher) for 30 min at 2.5 µM after resuspension in DMSO. Cells were treated with ROCK inhibitor Y27 (Y-27632 dihydrochloride, Y0503, Sigma) before wounding cells.

### Extracellular vesicles extraction

Extracellular vesicles were extracted by size exclusion chromatography. The medium was first concentrated by series of centrifugation using filters of 10 kDa (Centricon Plus-70 Ultracel-PL, 10 kDa cutoff-Millipore). The filter was washed with PBS at 3200g for 5 min. For a volume of 10 mL of KO medium, 4 mL of medium was added to the filter and centrifuge at 350 g at 4°C for 15 min. The supernatant was thrown away, 3 ml of KO medium was added to the filter for a centrifugation at 2000g for 20 minutes at 4°C. The supernatant was thrown away, and 3 mL of KO medium was added to the filter to be centrifuged at 3200g at 4°C for 25 minutes. A final volume of 500 µL was collected and stored at 4°C.

For the SEC, qEV-original-35nm columns (IZON) were used after 30 minutes at room temperature. The 500 µL of concentrated medium collected was placed at the top of the column. After the sample has completely entered the column, 2.5 mL of PBS was added to eluate the dead volume. 2.5 ml of PBS was added in the column to collect the first fraction of 500 µL containing EVs (Fraction 1); 2.5 mL of PBS was added to collect the intermediate fraction containing a small amount of EVs and soluble factors (Fraction 2); 2.5 mL was added to collect fraction 3 containing soluble factors and 2.5 mL of PBS was finally added to collect last fraction containing a very small amount of soluble factors (Fraction 4). The collected fractions were stored at -80°C before use.

### Image analysis and statistics

The intensity levels of pMLC and Vimentin in the immunofluorescence images were calculated by measuring mean fluorescent intensities of each cell using ImageJ.

The TFM images of the micropatterns were analyzed using a custom-designed macro in Fiji based on a previous study (Martiel et al., 2015). The topmost planes of beads before and after trypsinization were selected and aligned using a normalized cross-correlation algorithm (Align Slices in the Stack plug-in). The displacement field was computed from bead movements using particle image velocimetry (PIV). The parameters for the PIV analysis comprised three interrogation windows of 128, 64 and 32 pixels with a correlation of 0.60. Traction forces were calculated from the displacement field using Fourier transform traction cytometry and a Young’s modulus of 20 kPa, a regularization factor of 10−9 and a Poisson ratio of 0.5.

For migration assays, the nuclei of cells were manually tracked to determine the speed, directionality and persistence of migration. Mean velocity (𝑣), persistance (𝑝) and directionality (𝑑) of cell migration are calculated as follows: for a given (𝑥, 𝑦) coordinate of leading cell nucleus,

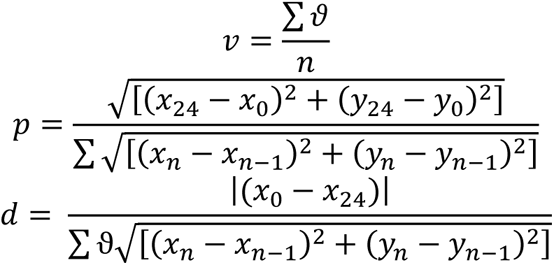

where 𝑛 is the number of time points acquired and 𝜗 is cell velocity.

For calculating the polarity index, images were processed using a custom ImageJ macro. First, fluorescent cell-adhesive patterns were individualized using a template matching method. Then nucleus and centrosome detection were realized based on threshold and size filtering within each cell of the doublet. Considering that only cell doublets were considered for further analysis, groups of cells encompassing two nuclei and two centrosomes were manually selected. The center of mass of the nucleus was computed and each centrosome was assigned to the closest nucleus. Finally, the nucleus-centrosome and nucleus-nucleus vectors were computed. Then, a post-processing step was carried out to quantify the orientation and connectivity of each cell within a doublet. The inter-nuclear distance was measured as the length of the nucleus-nucleus vector. The cell-cell connectivity was defined as the difference between the area of the convex envelope surrounding the cell doublet and the area of the cell doublet.

For calculating focal adhesion area, regions of interest were cropped and thresholded. The area of detected particles (adhesions) between 1 µm^2^ and 10 µm^2^ was plotted.

### Graphs and statistics

Statistical analysis was performed using GraphPad Prism software (Version 5.00, GraphPad Software). For each experiment, cell sampling and the number of independent replicates is indicated in figure legends. Data sets with normal distributions were analyzed with the One-Way Anova test to compare two conditions. Data sets were analyzed with a Mann-Whitney test. In the case of TFM and polarity assay measurements, outlier identification and data cleaning were performed due to apparent defects in some individual measurements. Results are presented as mean and SEM.

## Acknowledgements

This work was supported by the European Research Council: Consolidator Grant 771599 and Proof-of-Concept Grant 780458 to M.T., and an Advanced Grant 741773 (AAA) to L.B. J.G. was funded by CEA and Fondation ARC pour la recherche sur le cancer and is part of the Hematologie Oncogenese et Biotherapies (HOB) doctoral school. S.S. acknowledges support from Eric and Wendy Schmidt AI in Science Postdoctoral Fellowship, a Schmidt Futures Program. We thank Cédric Blanpain (Université Libre de Bruxelles) and Sebastien Jauliac (Hôpital Saint Louis) for scientific discussions during the initiation of the project. We also thank Clotilde Théry from the CurieCoreTech platform at Institut Curie for extraction of extracellular vesicles.

## Author contributions

J.G. and S.S. designed and performed the experiments, analysed and interpreted the results, and wrote the paper; B.V. assisted in the set-up and analysis of TFM and migration experiments, and helped with discussions; M.G. helped with image analysis; O. F. performed the CRISPR sequencing; L.B. and M.T. wrote the paper, helped with data interpretation and discussions, and supervised the project.

## Supplementary Information

**Supplementary Figure 1.**
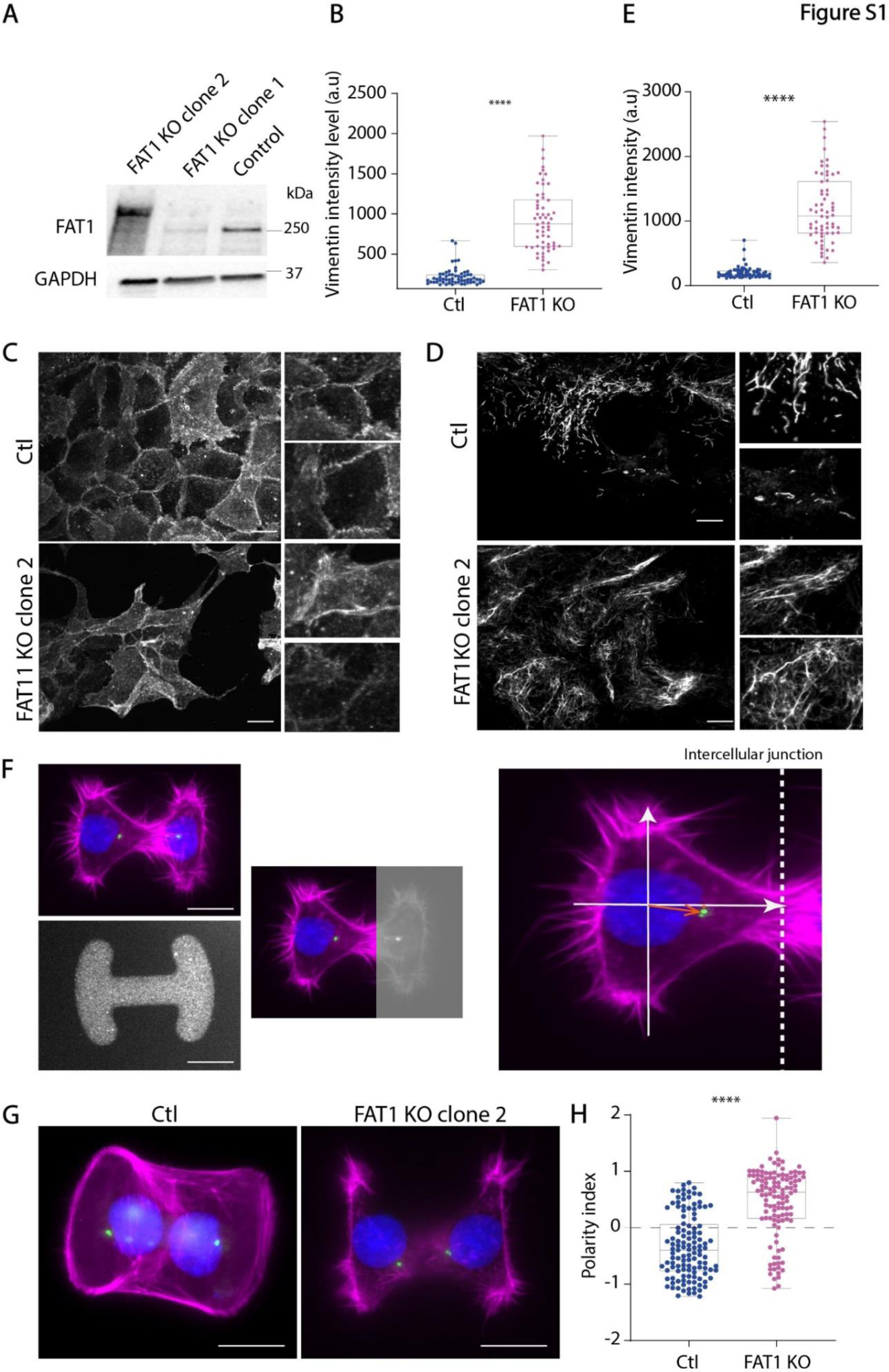
(A) Western-blot images of control cells and FAT1 KO cells. (B) Graph represents vimentin intensity in each condition. Represented data shown are from three independent experiments, for which n was between 10 and 20 cells for each cell type and conditions. (C) Confocal images of control and FAT1 KO clone 2 cells stained with E-cad, showing at a maximum intensity projection of 10 slices, spaced 1 micron and zoom images on the right. Scale bar: 10 µm. (D) Confocal images of control and FAT1 KO clone 2 cells stained with Vimentin, showing at a maximum intensity projection of 10 slices, spaced 1 micron and zoom images on the right. Scale bar: 10 µm. (E) Graph represents vimentin intensity in each condition. Represented data shown are from three independent experiments, for which n was between 10 and 20 cells for each cell type and conditions. (F) Confocal images of doublet of control or KO cells on H shaped micropatterns, size 1100 mm², stained for F-actin, pericentrin and DAPI, at a maximum intensity projection of 8 slices, spaced 1 micron. Scale bar: 20 µm. (G) Graph represents polarity index measurements. When 0.5<PI<2, centrosome is close to the cell-cell junction; when -0.5>PI>-2, centrosome is at the opposite side of cell-cell junction. Horizontal lines indicate the median value for each cell type. Represented data shown are from three independent experiments, for which n was between 20 and 30 cells for each cell type. (H) Representative images showing polarity index calculations. For all graphs, ****p < 0.0001 using Mann-Whitney T-Test.

**Supplementary Figure 2.**
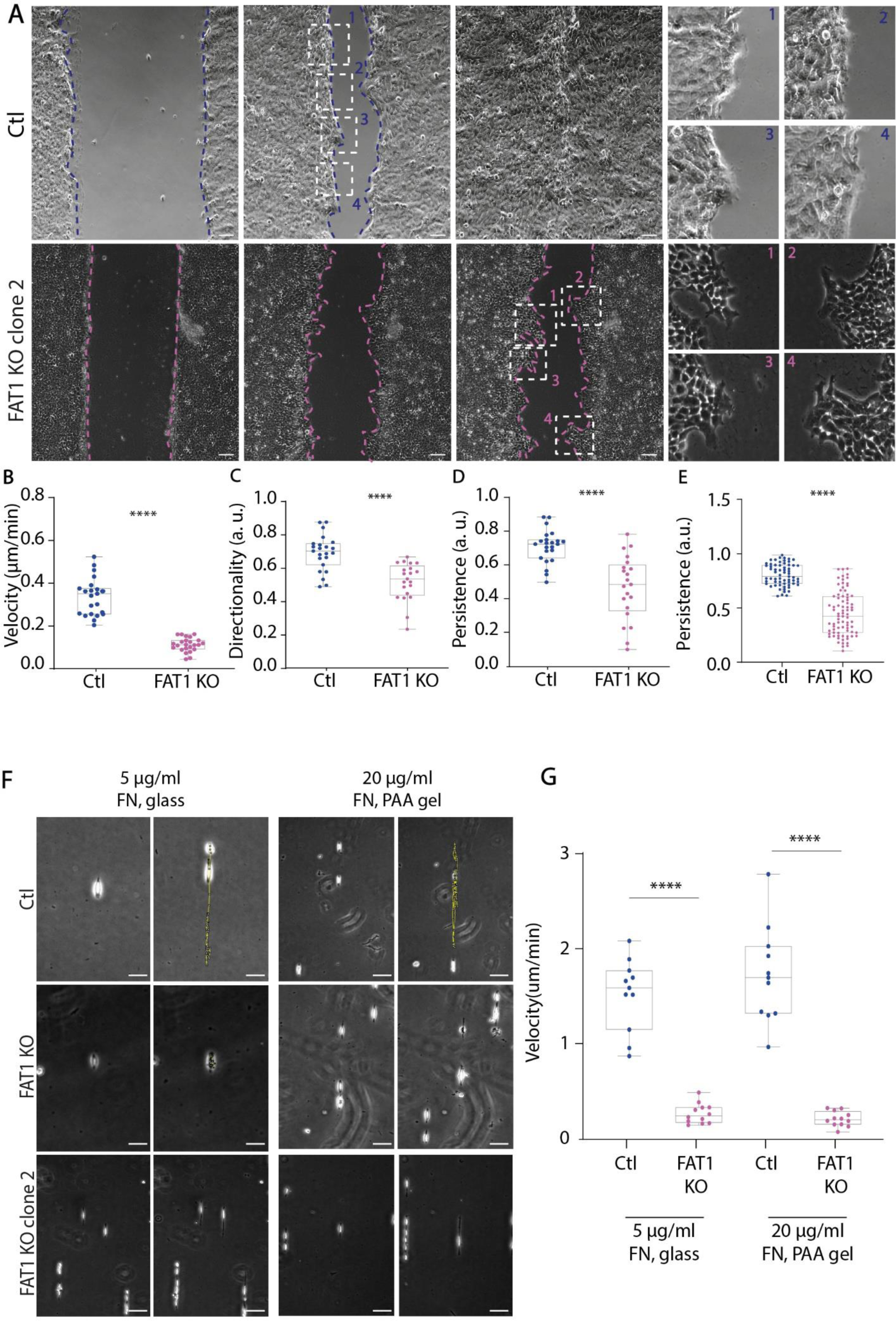
(A) Transmitted light images of control and KO clone 2 cells migrating at the indicated times after monolayer wounding. Images were acquired by time-lapse microscopy for 24 h, time interval 15 min. Scale bar: 100 μm. (B) Graph represents cell velocity from wound healing. Represented data shown are from three independent experiments, for which n was between 20 and 30 cells for each cell type. (C) Graphs represent cell directionality of cells. Represented data shown are from three independent experiments, for which n was between 30 and 50 cells for each cell type. (D, E) Graphs represent cell persistence of cells (FAT1 KO clone 1 and 2 respectively). Represented data shown are from three independent experiments, for which n was between 30 and 50 cells for each cell type. (F) Transmitted light images of single cell migrating control and KO on micropatterned lines, coated with Fibronectin at 5 µg/mL, or plated on 20 kPa gels coated with 20 µg/mL fibronectin. Dotted yellow lines show the trajectories of a single cell. (G) Graph represents single cell velocity. Plotted data come from three independent experiments, for which n was between 10 and 15 cells for each cell type. For all graphs, ****p < 0.0001 using Mann-Whitney T-Test.

**Supplementary Figure 3.**
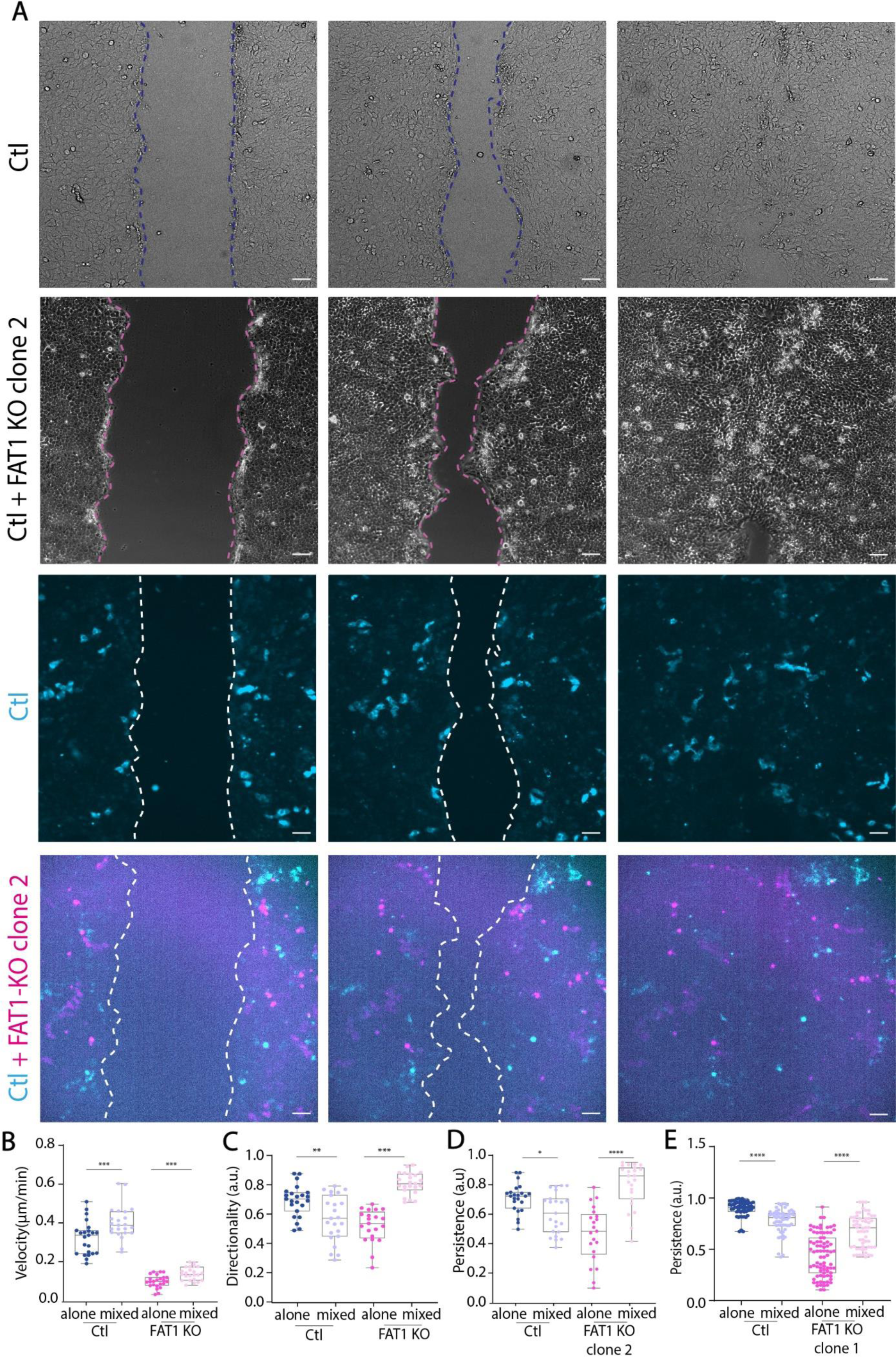
(A) Transmitted light and fluorescent images of control and mix population control+FAT1KO clone 2 at the indicated time after monolayer wounding. control cells were stained with Orange Cytotracker and are shown in blue, FAT1KO clone 2 cells were stained with Far-Red Cytotracker and are shown in pink. Images were acquired by time-lapse microscopy for 24 h, time points were taken every 15 min. Scale bar: 100 μm. (B) Graph represents cell velocity of migrating cells from wound healing. Represented data shown are from three independent experiments, for which n was between 40 and 50 cells for each cell type. (C) Graph represents directionality of cells. Represented data shown are from three independent experiments, for which n was between 20 and 30 cells for each cell type. (D) Graph represents persistence of cells. Represented data shown are from three independent experiments, for which n was between 20 and 30 cells for each cell type. (E) Graph represents persistence of cells of control and clone 1. Represented data shown are from three independent experiments, for which n was between 20 and 30 cells for each cell type. For all graphs, ****p < 0.0001 using Mann-Whitney T-Test.

**Supplementary Figure 4.**
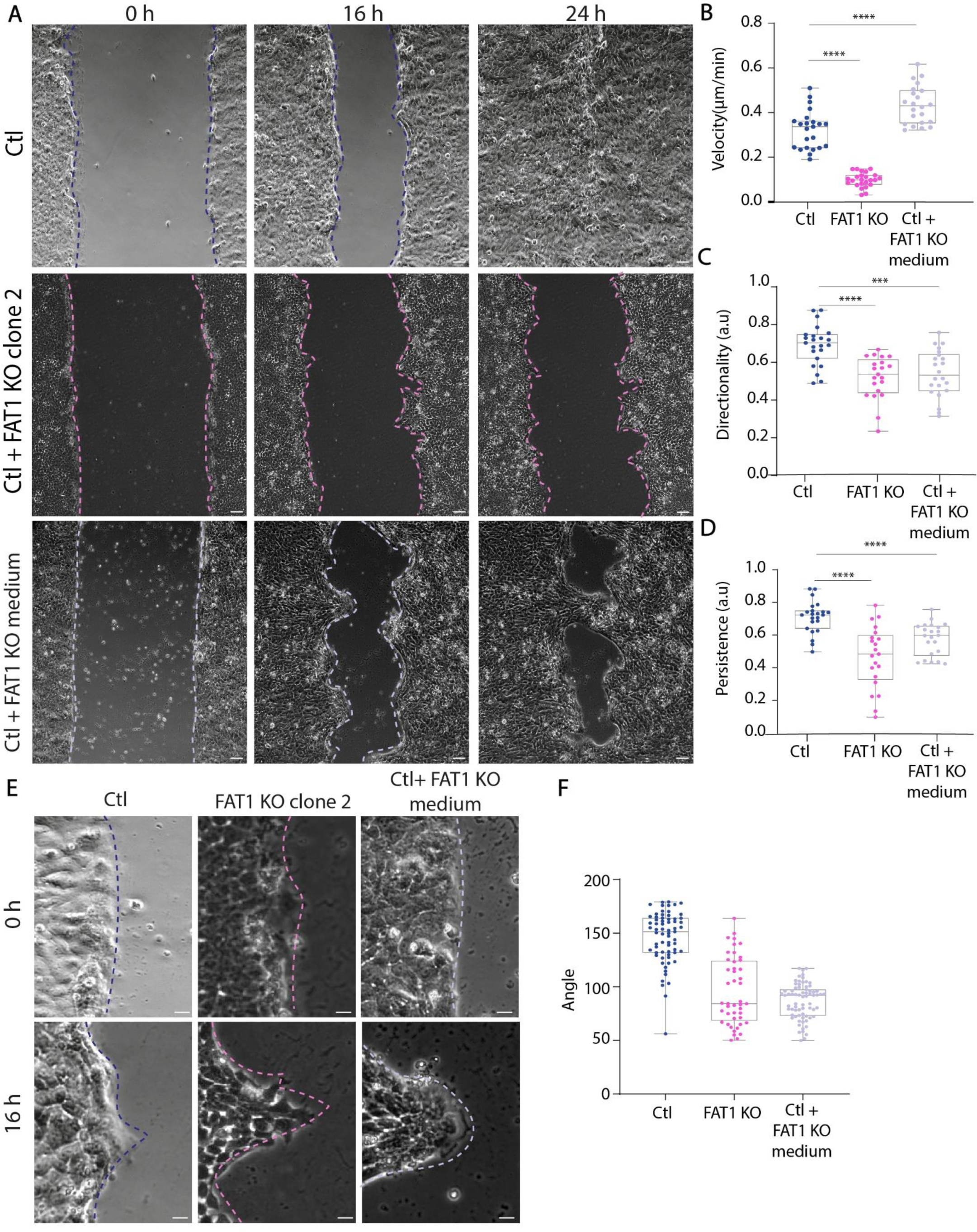
(A) Transmitted light images of control, FAT KO clone 2, control+ KO clone 2 medium cells migrating at the indicated times after monolayer wounding. control cells were treated with FAT1KO clone 2 medium one hour before wounding. Images were acquired by time-lapse microscopy for 24 h, time point 15 minutes. Scale bar: 100 μm. (B) Graph represents cell velocity of migrating cells from wound healing. Represented data shown are from three independent experiments, for which n was between 30 and 40 cells for each cell type and condition. (C) Graph represents cell directionality of cells. Represented data shown are from three independent experiments, for which n was between 30 and 40 cells for each cell type. (D) Graph represents persistence of cells. Represented data shown are from three independent experiments, for which n was between 20 and 30 cells for each cell type. (E) Representative images of angle measurements for each cell type and conditions. Angles measurements were obtained by the average of angle of the front every hour from wound healing movies. Scale bar: 10 μm. (F) Graph represents angle measurements for each cell type and conditions. Represented data are from three independent experiments, for which n was between 10 and 20 migrating fingers for each cell type and conditions. For all graphs, ****p < 0.0001 using Mann-Whitney T-Test.

**Supplementary Figure 5.**
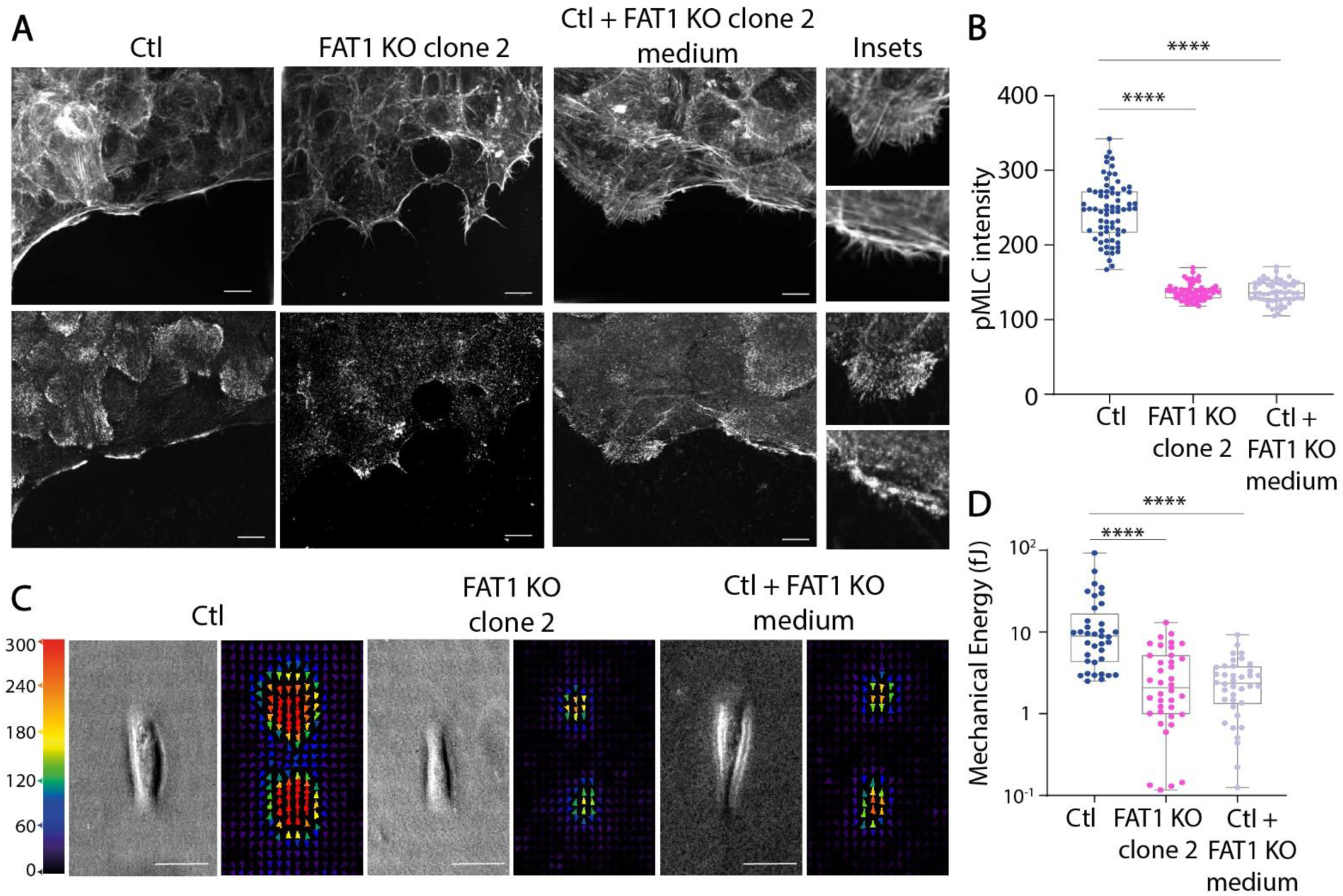
(A) Confocal images of cells stained with F-actin and pMLC staining in cells, showing a maximum intensity projection of 11 slices, spaced 1 micron, zoom images on the right for the control treated with FAT1KO medium. Scale bar: 10 μm. (B) Graph represents pMLC intensity in each condition. Represented data shown are from three independent experiments, for which n was between 20 and 30 cells for each cell type and conditions. (C) Stress-field maps of control or FAT1 KO clone 2 cells on rectangle-shaped micropatterned PAA gels of 20 kPa rigidity. (D) Graph represents Mechanical energy released by each cell type. Represented data shown are from three independent experiments, for which n was between 20 and 30 cells for each cell type and conditions. For all graphs, ****p < 0.0001 using Mann-Whitney T-Test.

**Supplementary Figure 6.**
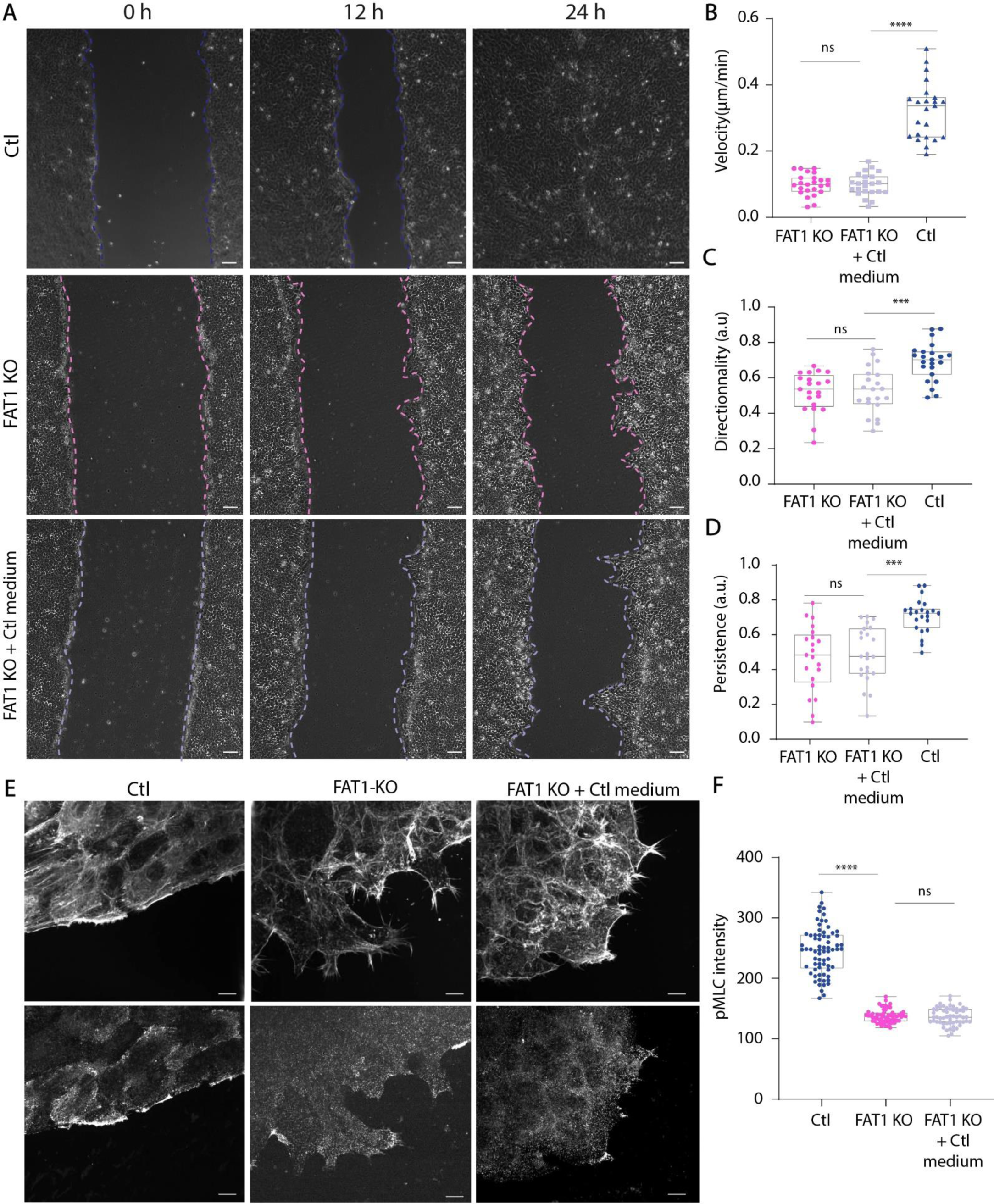
(A) Transmitted light images of control, FAT KO clone 1 and 2 cells migrating at the indicated times after monolayer wounding. FAT1 KO cells were treated with control medium one hour before wounding. Images were acquired by time-lapse microscopy for 24h, time point 15 minutes. Scale bar: 100 μm. (B) Graph represents cell velocity of migrating cells from wound healing. Represented data shown are from three independent experiments, for which n was between 30 and 40 cells for each cell type and condition. (C) Graph represents cell directionality of cells. Represented data shown are from three independent experiments, for which n was between 30 and 40 cells for each cell type. (D) Graph represents cell directionality of cells. Represented data shown are from three independent experiments, for which n was between 30 and 40 cells for each cell type. (E) Confocal images of cells stained with F-actin staining in cells and pMLC, showing a maximum intensity projection of 11 slices, spaced 1 micron, zoom images on the right for the KO cells treated with control medium. Scale bar: 10 μm. (F) Graph represents pMLC intensity in each condition. Represented data shown are from three independent experiments, for which n was between 20 and 30 cells for each cell type and conditions. For all graphs, ****p < 0.0001 using Mann-Whitney T-Test.

## Notes

### Competing Interest Statement

The authors have declared no competing interest.

